# Improving population scale statistical phasing with whole-genome sequencing data

**DOI:** 10.1101/2023.12.07.570528

**Authors:** Rick Wertenbroek, Robin J. Hofmeister, Ioannis Xenarios, Yann Thoma, Olivier Delaneau

## Abstract

Haplotype estimation, or phasing, has gained significant traction in large-scale projects due to its valuable contributions to population genetics, variant analysis, and the creation of reference panels for imputation and phasing of new samples. To scale with the growing number of samples, haplotype estimation methods designed for population scale rely on highly optimized statistical models to phase genotype data, and usually ignore read-level information. Statistical methods excel in resolving common variants, however, they still struggle at rare variants due to the lack of statistical information. In this study we introduce SAPPHIRE, a new method that leverages whole-genome sequencing data to enhance the precision of haplotype calls produced by statistical phasing. SAPPHIRE achieves this by refining haplotype estimates through the realignment of sequencing reads, particularly targeting low-confidence phase calls. Our findings demonstrate that SAPPHIRE significantly enhances the accuracy of haplotypes obtained from state of the art methods and also provides the subset of phase calls that are validated by sequencing reads. Finally, we show that our method scales to large data sets by its successful application to the extensive 3.6 Petabytes of sequencing data of the last UK Biobank 200,031 sample release.

## Introduction

In the era of biobanks, population-scale sequencing is becoming increasingly common [1], producing variant calls and comprehensive data sets that depict the genomic landscape across a vast number of samples [2]. To enable analyses at the haplotype level, it is necessary to phase the genotype data produced by sequencing. When dealing with data sets comprising thousands to millions of samples, statistical methods like SHAPEIT5 [3] or Beagle5 [4] are the standard approach. These methods borrow information across many samples in the population in order to produce precise haplotype estimates for common variants. Nevertheless, they tend to exhibit higher error rates when handling rare variants due to the limited information available for those in the population. Conversely, haplotype assembly methods, like WhatsHap [5] for instance, utilize local read reassembly techniques to group nearby variants into fully resolved haplotype blocks, often called phase sets. This type of approach offers the advantage of phasing variants with an accuracy level that remains independent of allele frequency, but typically provides only partial estimates, often spanning a few kilobases at most [5]. Indeed, variants located too far apart cannot be linked by the same set of reads and thus cannot be phased together. To harness the strengths of both approaches, a good strategy is to combine statistical phasing methods with haplotype assembly methods, as previously proposed in the phasing pipeline based on WhatsHap [5] and SHAPEIT4 [6]. While this approach leads to high accuracy levels, it faces limitations in scaling to the vast amounts of samples present in modern sequencing data sets, comprising thousands or even millions of samples.

In this paper, we address this challenge and introduce an accurate and efficient haplotype alignment method, the Smart and Accurate Polishing of Phased Haplotypes Integrating Read Enhancements (SAPPHIRE) method. SAPPHIRE is primarily designed to refine the haplotypes estimated by SHAPEIT5 [3]. Our novel method capitalizes on the phasing confidence scores provided by SHAPEIT5 to pinpoint poorly phased rare variants. Then, it employs local read realignment techniques to correct errors at these poorly phased sites. This targeted approach leads to a substantial reduction in the volume of sequencing data that requires processing, reducing it by orders of magnitude. This makes it a viable solution even for extraordinarily large sequencing data sets. To illustrate its speed and accuracy, we applied it to an extensive data set comprising 200,031 UK Biobank samples, each with whole-genome high-coverage sequencing, totaling more than 3.5 Petabytes of data. Our findings show that our method can traverse all this sequencing data at a relatively modest computational cost and achieve a significant enhancement of statistically estimated haplotypes, notably at rare variants and singletons.

## Results

### Overview of the SAPPHIRE method

Similar to long-read polishing [7–9], where errors in long-reads are corrected by aligning highly accurate short-reads, SAPPHIRE corrects phase errors in estimated haplotypes by aligning highly accurate short reads. The type of error introduced by statistical phasing can be classified into two categories, often dependent on allele frequency (see Fig 1A). At common variants, we encounter switch errors, where entire contiguous segments of haplotypes are incorrectly phased. At rare variants, we encounter flip errors, where only a single heterozygous genotype is flipped onto a correctly phased haplotype background. In large-scale sequencing data sets, such as the UK Biobank, the large number of samples being sequenced implies (i) very good haplotype estimates at common variants and thus few switch errors and (ii) an excess of rare variants which makes flip errors the primary source of error in this type of data sets. In this work, we propose an approach to quickly identify flip errors in large scale sequencing studies, and to correct these errors by performing local re-alignment of the overlapping sequencing reads (flip error detection and correction, Fig 1B). To achieve this, we first employ SHAPEIT5 for statistical phasing, as it provides a phasing confidence score for each rare heterozygous genotype. This score essentially corresponds to the probability of the phase the software reports and enables us to pinpoint the subset of rare heterozygous genotypes with low phasing confidence. Then, we extract sequencing reads overlapping these poorly phased variants and use these reads to re-phase all these problematic variants, when possible. In practice, we consider each of these variants in turn and check if any of the extracted reads can connect the variant of interest to a nearby common heterozygous variant. By checking the allelic content of these phase-informative reads, we can collect evidence (number of reads) that confirm (validate) or invalidate the phase relationship reported by statistical phasing. Each time the phase relationship between a low confidence-phased heterozygous genotype to its neighbors is invalidated, we flip it. For high-confidence genotypes (including common variants), we do not make any adjustments, and assume they are correctly phased. With this method, even singletons (minor allele count of 1 in the population) can be phased if they are within read-pair distance of another nearby common variant. It is worth noting that this approach can also be used to refine the output of other statistical phasing methods, even though they do not provide phasing confidence scores. In such cases, we propose using only allele frequency as a proxy for phasing confidence to identify possible phasing errors.

**Fig 1.**
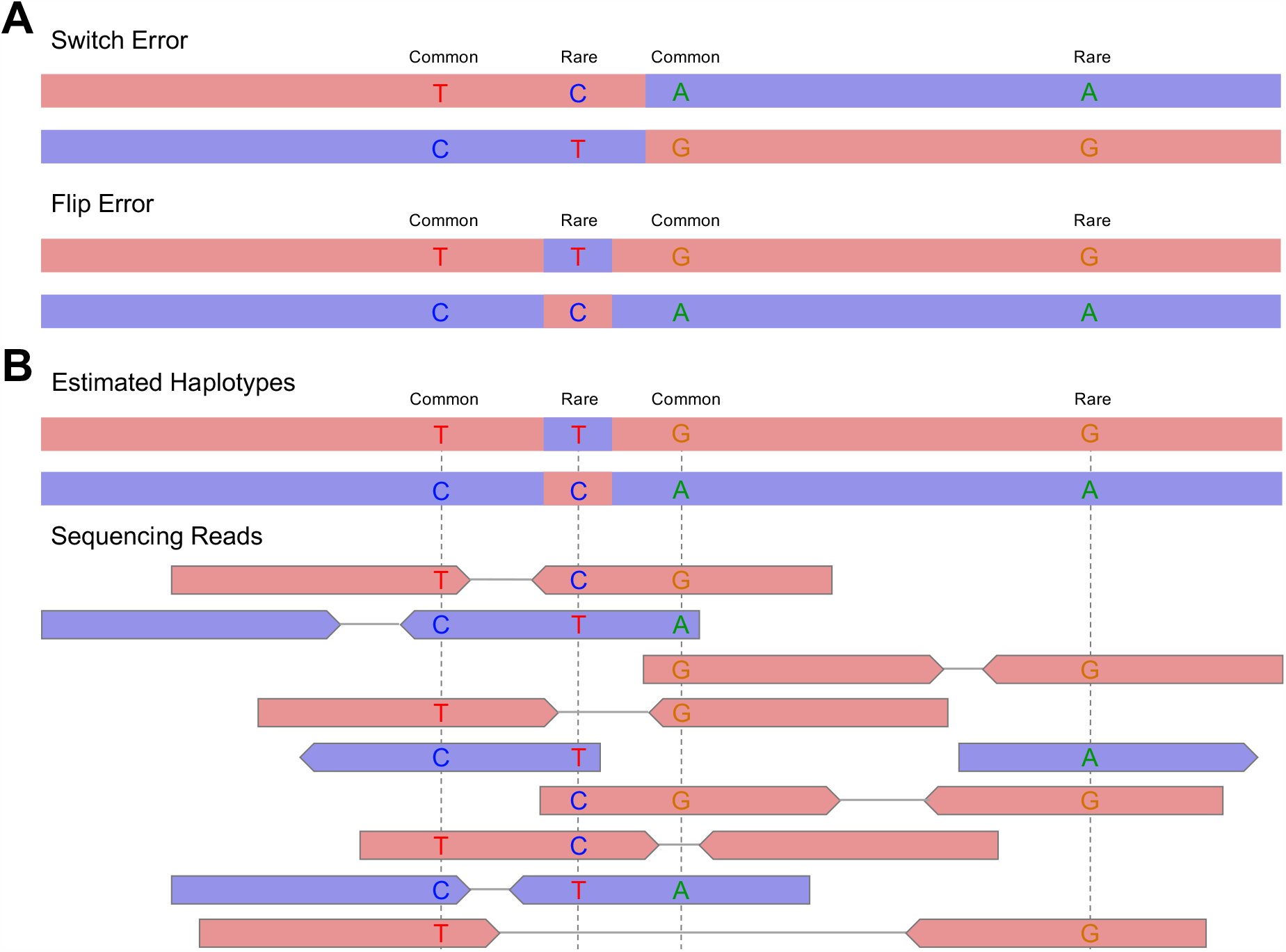
Phasing errors and read-based validation of phase. **A**. Phasing errors come in two types, switch errors where the entire contiguous segments of haplotypes are phased incorrectly and flip errors where only a single heterozygous genotype is flipped on a correctly phased haplotype background. **B**. Whole-genome sequencing reads aligned to the estimated haplotypes allow to validate or contradict the phase relationship of heterozygous variants that are in close enough proximity to be carried by a single sequencing read or a pair of sequencing reads. In this example we can see that the two common heterozygous variants have been phased correctly with regards to each other. However, the rare variant T/C was phased incorrectly (assigned to the wrong haplotype) and should therefore be switched. The rare variant G/A is phased correctly.

### SAPPHIRE on the UK biobank with whole-genome sequencing data

We applied the SAPPHIRE method described above on two data sets of the UK Biobank. (1) Wholegenome sequencing (WGS) data on chromosome 20 for 147,754 samples, a subset of the 150,119 samples release [10] with parental genomes of duos and trios excluded for validation. SAPPHIRE was applied on this data set, after a first statistical phasing pass through SHAPEIT5 as presented in [3]. (2) WGS data for all autosomes of the newest release of 200,031 samples also phased with SHAPEIT5 [11]. The data set of (1) was used to estimate the phasing accuracy using the offspring genomes as validation data. The data set of (2) includes the full set of samples (parental and offspring genomes) and was used to compute the costs of the SAPPHIRE method at scale, when applied genome-wide to WGS data for a large number of samples. In addition, we also used (2) to study the parental origin of de novo mutations, evaluate the number of genotypes that could be reassessed, and retrospectively evaluate the phasing confidence scores provided by SHAPEIT5.

### Phase quality of UK Biobank data after SAPPHIRE

The 147,754 sample data set was used to assess the improvement of phasing quality our method could lead to when applied on the haplotypes produced with SHAPEIT5 [3]. We used parental and offspring genomes as validation (31 trios, 432 duos) to 1) derive a true set of haplotypes for the offspring using inheritance logic 2) to compare the offspring haplotypes obtained by statistical phasing methods SHAPEIT5 [3], Beagle5.4 [12], and the SAPPHIRE method, to the true set of 1). The phasing quality is measured by computing the switch error rate (SER), which is the fraction of successive heterozygous genotypes phased incorrectly as defined in [3]. Because statistical phasing methods have a switch error rate that is correlated with the frequency of a variant, we measured switch error rates within bins of minor allele counts (MAC). Phasing performance is reported in Fig 2 and shows that the polished haplotypes exhibit lower phasing error rates at all minor allele count bins for rare variants. This is particularly apparent for singletons, where we go from 34% before SAPPHIRE to 21% after, a reduction of 13% in error rates. Importantly, these results include all singletons, covered or not by phase informative reads. When focusing on the subset of genotypes that can be re-phased using our approach, which comprises 43% of all singletons, the error rate drops to 2.9% (Fig 2). Overall, the results demonstrate that our method substantially improves the phasing quality at variants with a minor allele count below 200 (minor allele frequency (MAF) *<* 0.27%), with the highest boost observed at singletons.

**Fig 2.**
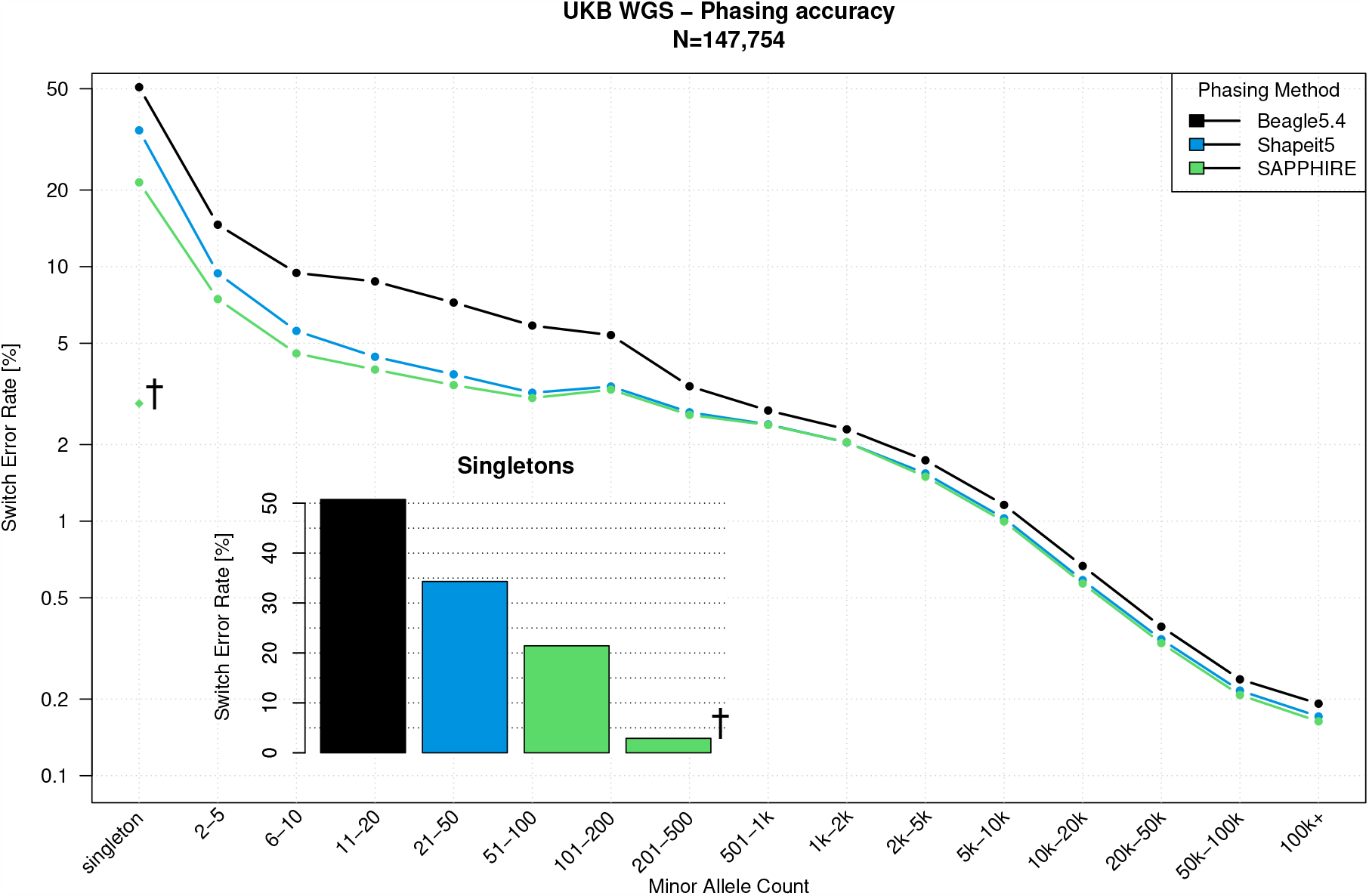
Phasing performance. Switch error rate (SER, y-axis, log-scale) of the polished haplotypes with SAPPHIRE (green) compared to SHAPEIT5 (blue) and Beagle5.4 (black) stratified by minor allele count (x-axis) for the UK Biobank whole-genome sequencing chromosome 20 data. The † shows the switch error rate (2.9%) for singletons that have been rephased by the SAPPHIRE method (43% of all singletons)

### Assessment of the phasing quality of de novo mutations

To biologically validate the haplotypes and to showcase a possible downstream usage, we looked at de novo mutations (DNMs). DNMs are mutations that are not present in the parental genomes. They can occur in an egg or sperm cell of a parent, in the fertilized egg after the egg and sperm unite, or in another type of cell division during embryo development [13]. Note that only a minority of DNMs are postzygotic as shown in a recent study assessing 6,034 DNMs in 91 monozygotic twins in which 97.1% of the DNMs were found in both twins [14] (and therefore are not postzygotic). DNMs are true singletons even in trio data sets as they are new mutations and do not appear in the parental genomes. However, these DNMs still require to be phased to either the paternal or maternal haplotype, which is impossible to achieve using parental genomes. All non postzygotic (most DNMs) have a parental origin that can be deduced from the haplotype on which they lie. Multiple studies have shown that DNMs originate more often from the paternal side than from the maternal side, with a ratio of between 3 : 1 and 4 : 1 [14–17]. In our work, we evaluated the phase of 3,744 SNV singletons (de novo mutations) in the offspring of 93 trios from the UK Biobank whole-genome sequencing data. Out of the 3,744 DNMs, 1,689 (45%) could be linked to a paternal or maternal haplotype using sequencing reads. Fig 3 summarizes the phase identification of DNMs. The phased DNMs show a total mean 1, 271 : 415 ≈ 3.06 : 1 paternal/maternal ratio and a median ratio of 14 : 4 = 3.5 : 1 paternal/maternal DNMs which is in line with previous findings. If we use the phase given to these 1,689 DNMs by SHAPEIT5 the total mean ratio is 1096 : 590 ≈ 1.86 : 1 and median ratio of 12 : 6 = 2 paternal/maternal DNMs which is further away from the expected result because of phasing errors. It has also been shown that the number of DNMs in offspring is positively correlated with increasing paternal age at the time of conception [14–17]. The data from the 93 trios was used to assess this relation. Due to the age distribution within the UK Biobank (40-69 years), the age range of parents in trios is limited to young adults. However, results shown in Fig 4 still exhibit a positive correlation with the age of the father at conception, although not significant (*p*-value = 0.318). And as expected no correlation with the age of the mother (*p*-value = 0.926). The replications of these patterns demonstrates that the phasing of the SAPPHIRE method at singletons, variants at which statistical phasing performs very poorly, is of high quality and can be used to reveal important biological properties.

**Fig 3.**
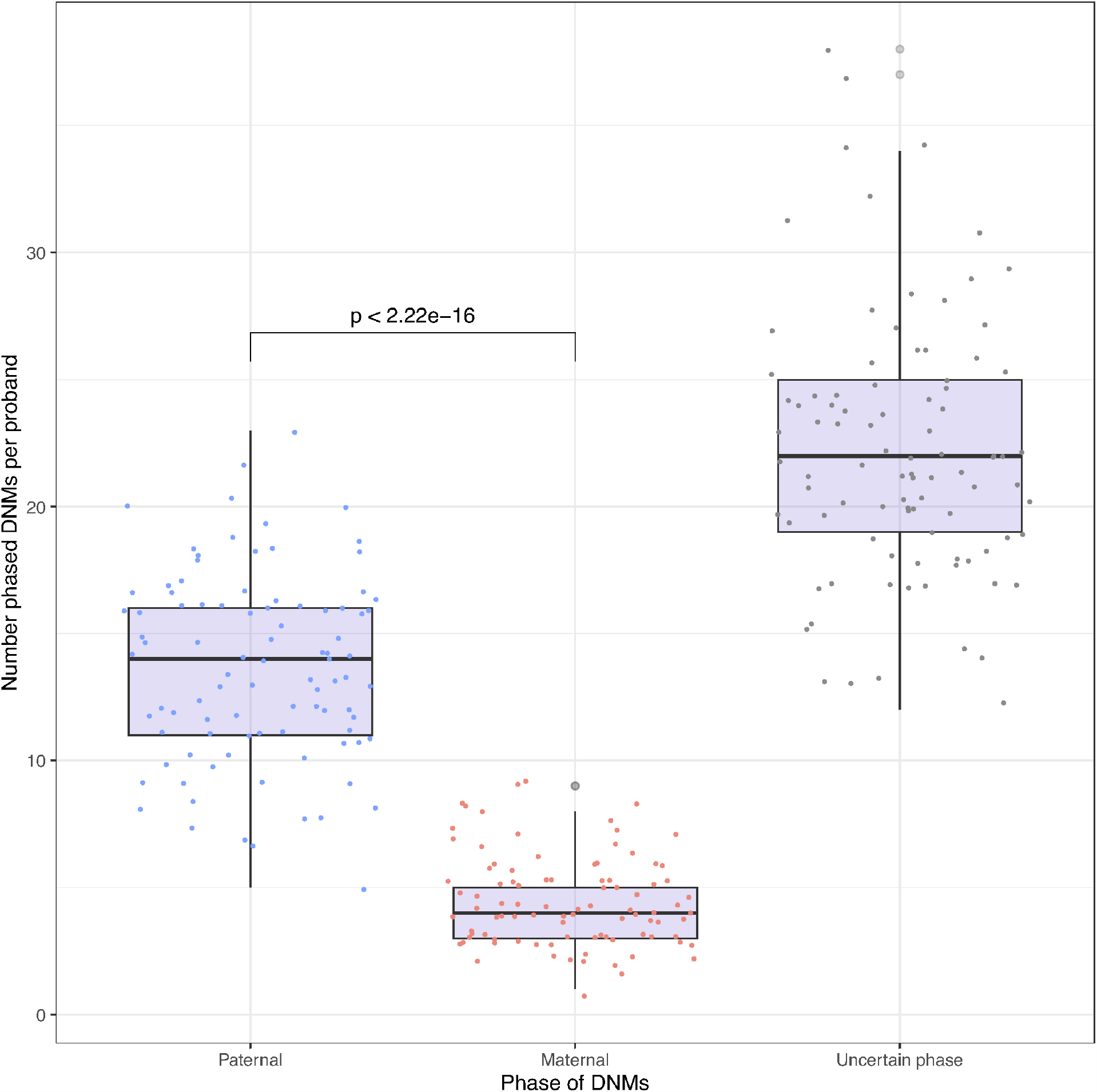
Distribution of de novo mutations (DNMs) either on the paternal or maternal haplotype confirmed by WGS reads. DNMs or uncertain phase are DNMs that have been phased with low confidence and have no WGS reads to validate their phase. The paternal/maternal median ratio is 14 : 4 = 3.5 : 1 paternal/maternal DNMs. *p*-value between paternal and maternal categories correspond to Wilcoxon test *p*-value

**Fig 4.**
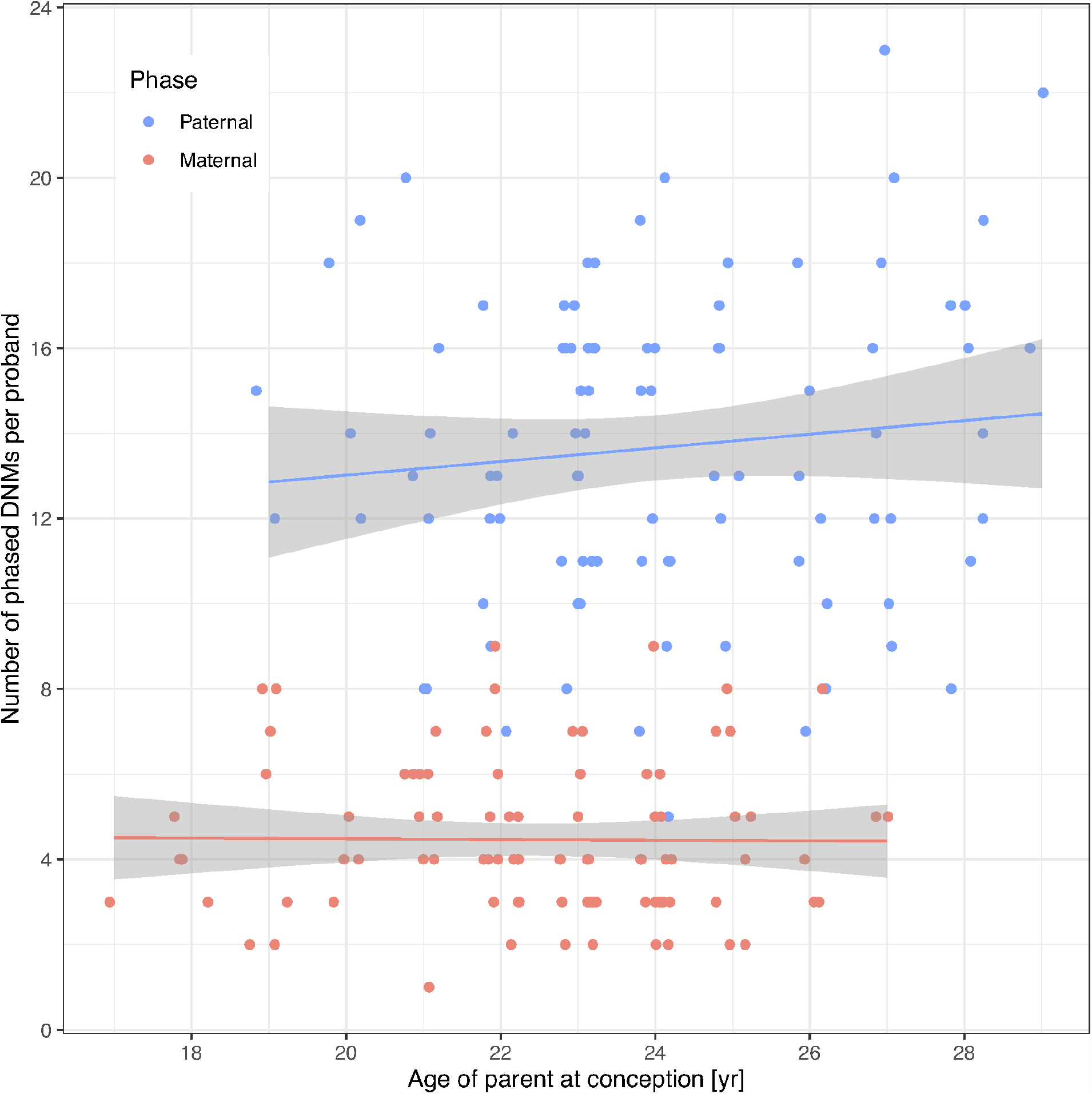
Number of de novo mutations (DNMs) per proband grouped by parent of origin relative to the age of parent at conception. Only DNMs with phase confirmed by WGS reads are plotted. A positive trend is observed relative to the age of the father at conception. No correlation with age is observed on the maternal side. *N* = 93.

### Number of rare variants that can be rephased through SAPPHIRE

Rephasing a rare variant requires close enough proximity to a common variant in order for both to be covered by the same sequencing reads or read pairs, as shown in Fig 1B. We therefore evaluated rare variants on chromosome 21 for 200,031 samples to see how many could be rephased using sequencing reads. We extracted a total of 11,471,867 heterozygous variants with a SHAPEIT5 confidence score below 0.99 (considered here as poorly phased with statistical phasing). Fig 5 shows the distribution of variants. Only SNVs (89.4%) were considered. In theory, other types of variants (e.g. short indels) can also be rephased but special care to the realignment model is needed to accommodate such cases. In total, 45.2% of the SNVs shared at least one high quality sequencing read or read-pair with a nearby common and accurately phased variant. Out of those, 69.7% were phased correctly by SHAPEIT5, while 30.3% required the original phase to be corrected. Finally, Fig 6 and 7 show the distribution of phase corrections per minor allele count (MAC) bin, as counts and percentages, respectively. As expected, the singletons require more correction, as these are not possible to phase reliably through statistical models.

**Fig 5.**
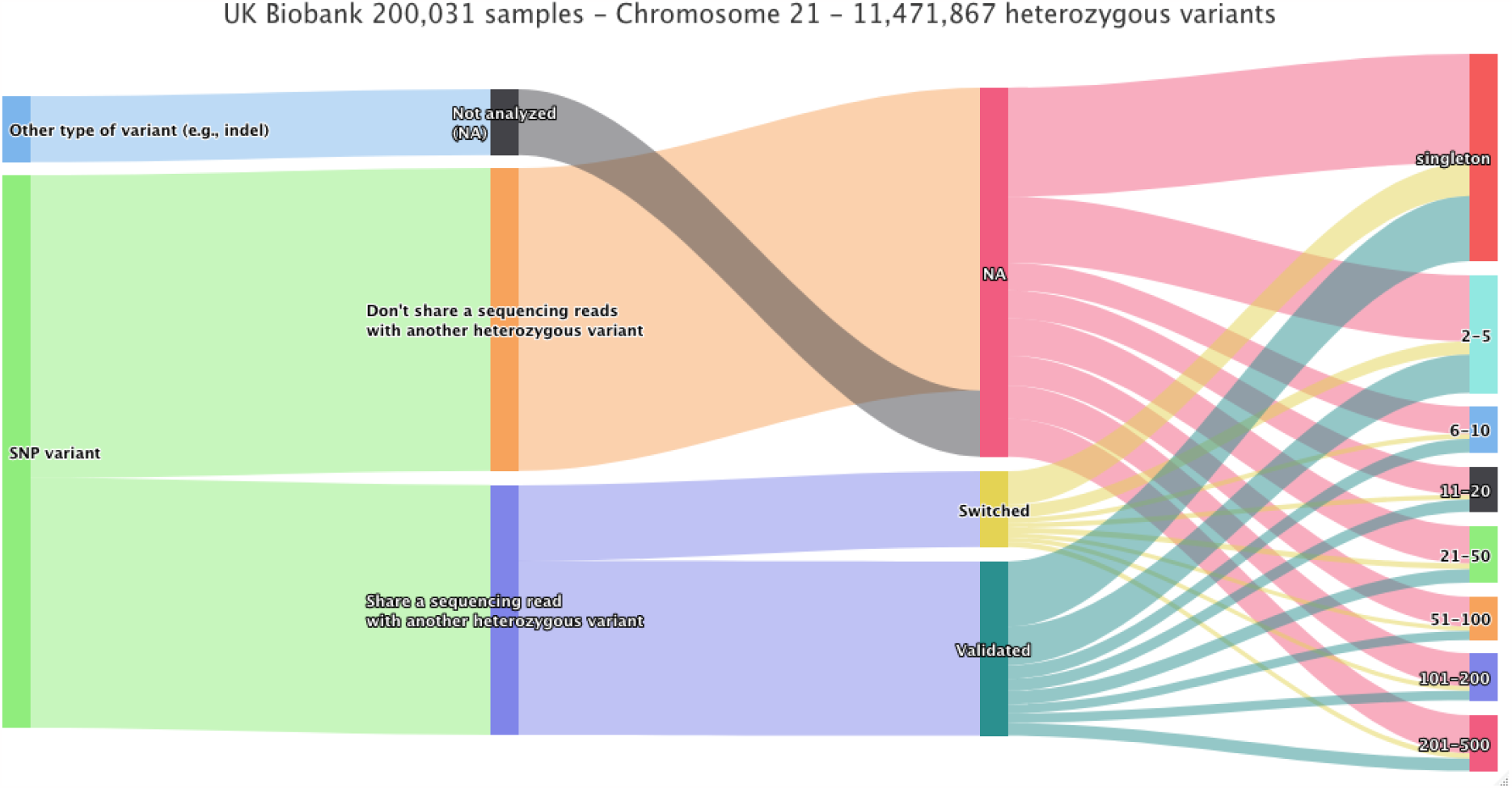
Distribution of heterozygous genotypes phased by SHAPEIT5 with phasing probability (PP) score less than 0.99. For the 200,031 sample release of the UK Biobank whole-genome sequencing calls, 11,471,867 heterozygous genotypes on chromosome 21 were extracted based on the phasing probability (PP) score given by SHAPEIT5. Out of all the genotypes with a PP score less than 0.99 only SNVs (89.4%) were considered for analysis, out of all SNVs 45.2% share at least one sequencing read or read-pair with a confidently phased genotype. Of all analyzed SNVs, 30.3% were phased incorrectly with regards to the other genotype on the reads. The last column displays the ratios of genotypes that were switched, validated, or not analyzed per minor allele count bin.

**Fig 6.**
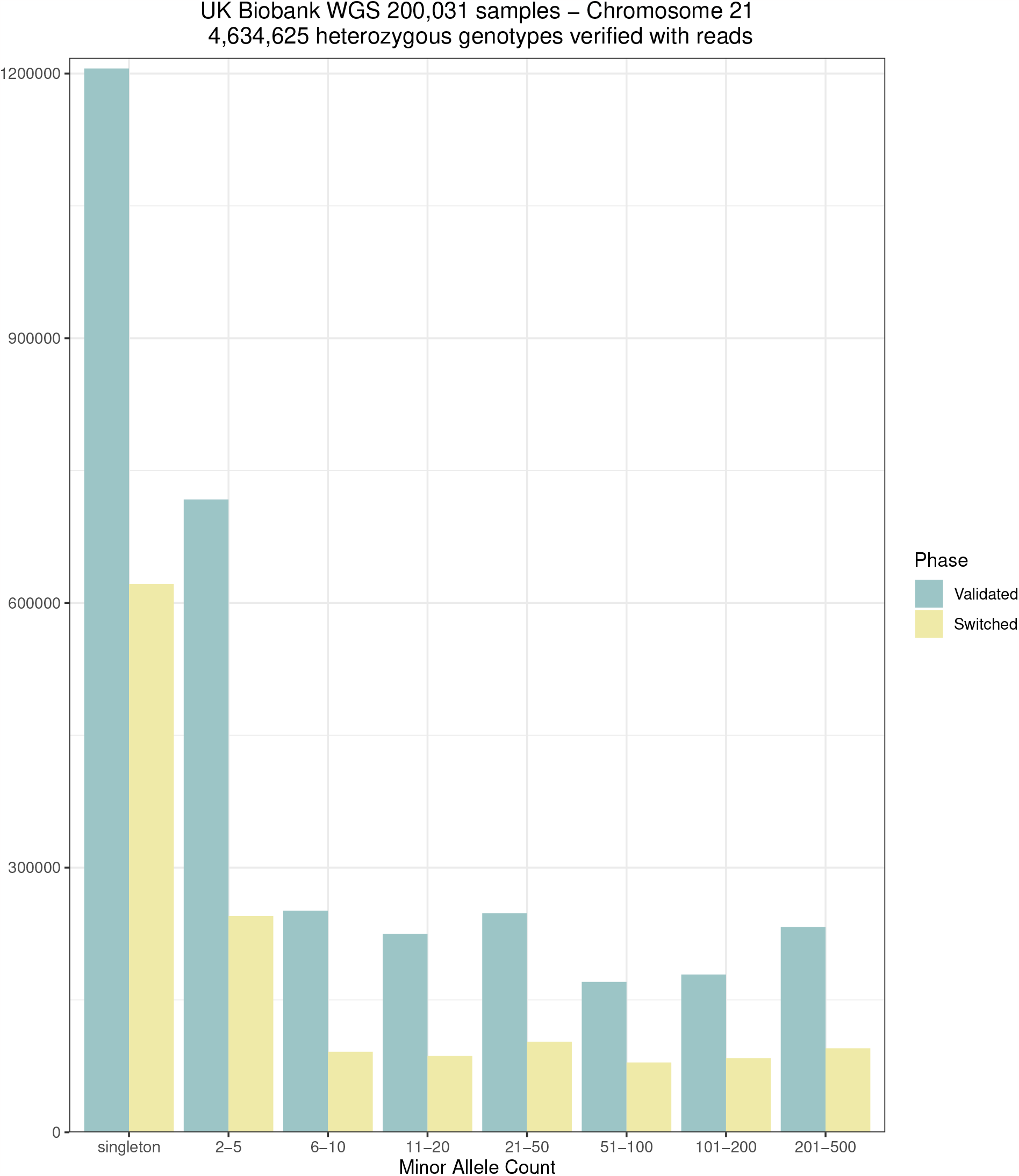
Phase validation. Number of heterozygous genotypes with the phase validated or switched grouped by minor allele count in the 200,031 samples data set on chromosome 21 when applying SAPPHIRE after SHAPEIT5.

**Fig 7.**
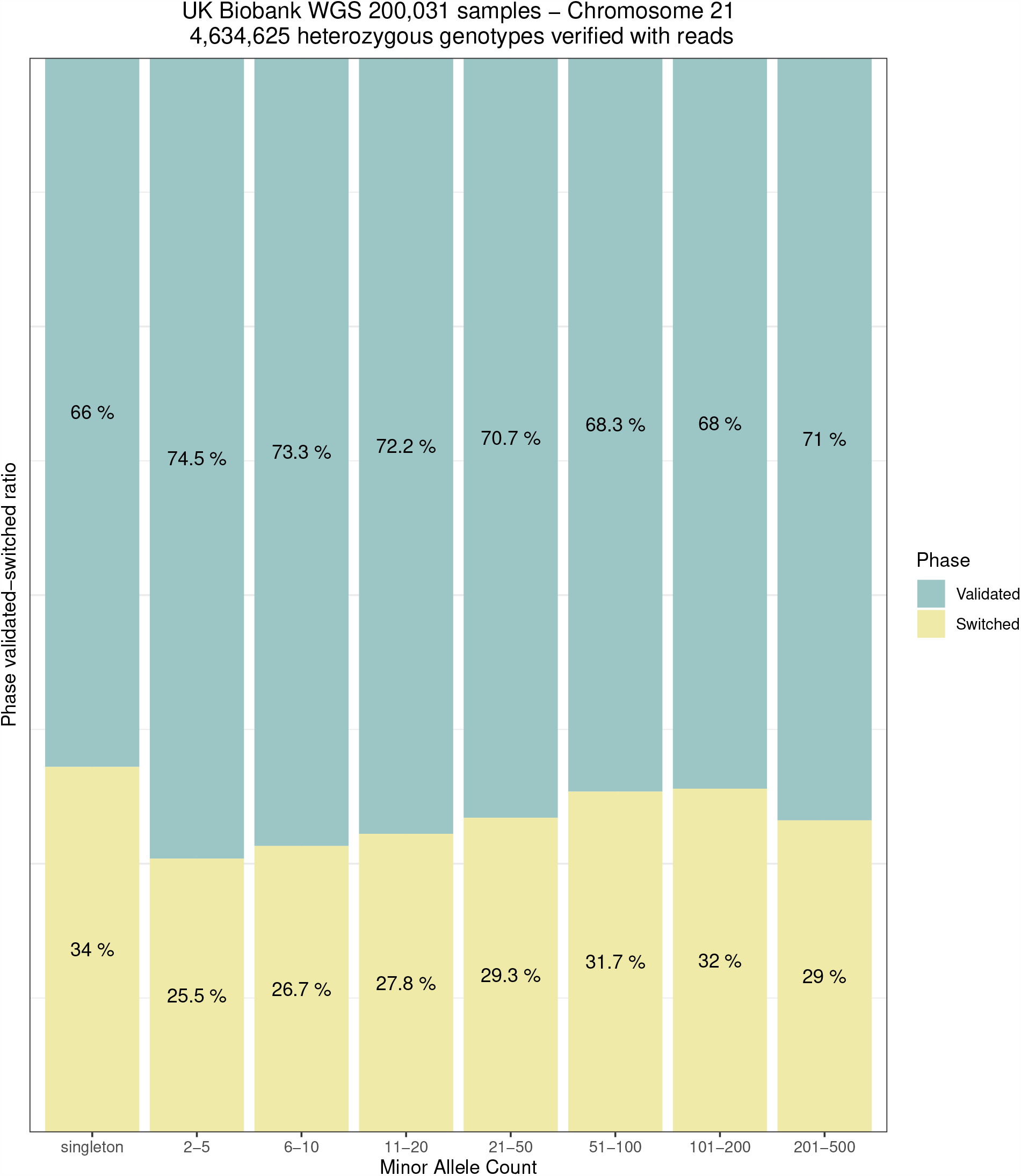
Phase validation distribution. Distribution of validated/switched phase calls of 4,634,625 genotypes grouped by minor allele count in the 200,031 samples data set chromosome 21 when applying SAPPHIRE after SHAPEIT5.

### Cost of the SAPPHIRE method

All computations have been performed on the Research Analysis Platform (RAP) of the UK Biobank. The cost to phase 147,754 samples on chromosome 20 with SHAPEIT5 was £74.9 [3]. In contrast, phase polishing the resulting haplotypes with SAPPHIRE cost only £5.0 which is less than 10% of the original cost. Phase polishing the full set of 200,031 samples of the UK Biobank across all chromosomes requires going through 200,031 CRAM files (on average 18 GB per sample), representing over 3.5 Petabytes (3,669,126 Gigabytes) of data in total. The total cost for this step was £570 (details given in the methods section). We also estimated the cost of running WhatsHap [5] to extract phase sets on the same dataset would add up to at least £50,000 (see methods). This demonstrates the need for a targeted approach compared to first extracting phase sets for all heterozygous variants from sequencing reads.

### Assessment of SHAPEIT5 phasing confidence

The confidence score given by SHAPEIT5 is a probability that ranges between 0.5 and 1.0. A score of 0.5 means full uncertainty in the phasing call (equivalent to a coin toss) while 1.0 means that the software is pretty certain on the call it reports. The SAPPHIRE method allows us to verify, based on sequencing reads, many of the original phase calls and we can collect statistics about their correctness. In practice, we binned the SHAPEIT5 phase calls by confidence scores (PP field) and computed the number of calls that were validated or invalidated by sequencing reads. We analyzed chromosome 21 for 200,031 samples and collected statistics for 4,634,625 phase calls. Fig 8 reports the percentages of phase calls that we validated or corrected using sequencing reads. Higher confidence scores come with lower error rates, ranging from 33.7% to 12.5%. For a phase confidence of 0.5, SHAPEIT5 performs better than a coin toss with only one third of the calls being incorrect. For a phase confidence score between 0.9 and 0.99, SHAPEIT5 seems a little overconfident. This shows that the SAPPHIRE method can also be used to evaluate and recalibrate phasing model confidence scores.

**Fig 8.**
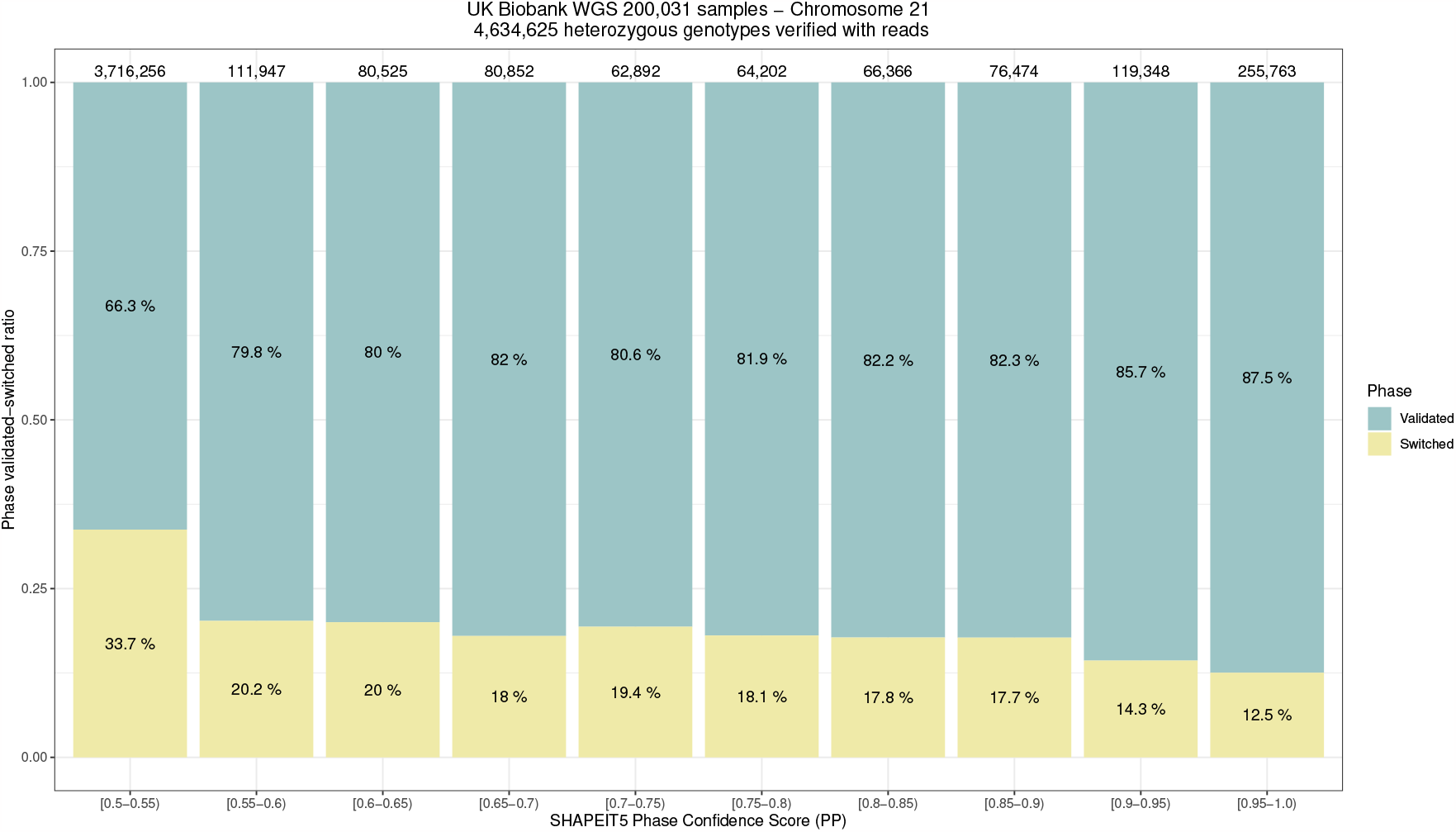
Assessment of SHAPEIT5 phasing confidence score (< 0.99) with sequencing reads. We evaluated the phasing confidence scores smaller than 99% given by SHAPEIT5 on its phase calls with sequencing reads. Out of 11,471,867 heterozygous genotypes on chromosome 21 for 200,031 samples we were able to check the phase of 4,634,625 with sequencing reads. The ratio of validated/switched phase calls is reported per phase confidence score bins as given by SHAPEIT5.

## Discussion

We present SAPPHIRE a novel approach to improve the phasing accuracy at rare variants within population-scale datasets initially phased with statistical methods. We show that we can reduce switch error rates for variants with low-occurrence frequencies with the most notable impact at extremely rare variants and singletons. Our method also delimits the subset of phased alleles that have been validated by sequencing reads, which makes it possible to discern them from statistically phased variants. We show that our method is scalable and can be used on Petabytes of sequencing data thanks to a targeted approach where only alleles of low confidence are analyzed. Phase polishing of the UKB WGS dataset with 200,031 samples phased with SHAPEIT5 through SAPPHIRE was achieved with a compute cost of £570, or less than £0.003 per sample. The rephased dataset shows reduced error rates on all rare variant frequency bins with major improvements on the phasing of singletons. SAPPHIRE not only improves overall estimated haplotype quality but can also be used as a benchmark for phasing in general as it can report the number of discordances between estimated haplotypes and phase relationships observed within sequencing reads.

The subset of read validated phased rare variants allows for new downstream analyses which were not possible with the lower accuracy of statistical methods, especially for singletons. To demonstrate the impact of accurately phasing singletons we replicated two key observations of de novo mutations: (i) we observed the expected paternal/maternal imbalance in the origin of the de novo mutations, and (ii) we found a positive correlation with the father’s age at conception and the number of de novo mutation in the proband. These observations could not be replicated through statistical phasing alone. We predict that our method will allow for further analyses of rare variation such as discovery of compound heterozygous gene knockouts.

We show that by combining statistical methods and targeted read based methods, we can provide highly accurate haplotype estimates at a low cost. The method can be applied on a per sample basis, on a sample subset of interest or on a whole population. If long read sequencing data becomes available for the samples, the method would allow linking even more variants and further improve the estimated haplotypes.

We believe that phase polishing large statistically phased data sets is the most effective method to date. By combining the effectiveness of statistical methods for common variants and the precision of reads for rare variants we achieve the highest possible accuracy. The order of the methods is of crucial importance, because of the high cost of processing large amounts of sequencing data, it is best to only target sites of low confidence given by the statistical phasing method (or of low minor allele frequency by proxy) in order to avoid spending time on sequencing data where it unnecessary. We estimated that the SAPPHIRE method has a cost at least a hundred times lower than current read-based methods and have shown that it is possible to polish all autosomes for 200,031 samples for a total of £570 improving the overall phasing quality of the data set especially at rare variants and even singletons.

## Methods

Phase polishing through the SAPPHIRE method is as follows: First, heterozygous genotypes from all samples are extracted. Then, the phase for each pair of genotypes with overlapping sequencing reads is verified, with common variants as reference for phase verification. Rare variants are checked against common ones. If sequencing reads clearly show a reversed phase, SAPPHIRE corrects it and the read count supporting the phase is reported. Unchanged variants also have their read count reported for phase call confidence. Finally, all switched or validated genotypes are updated in the original VCF file to include the new updated phase and supporting read count. The pipeline is depicted in Fig 9.

**Fig 9.**
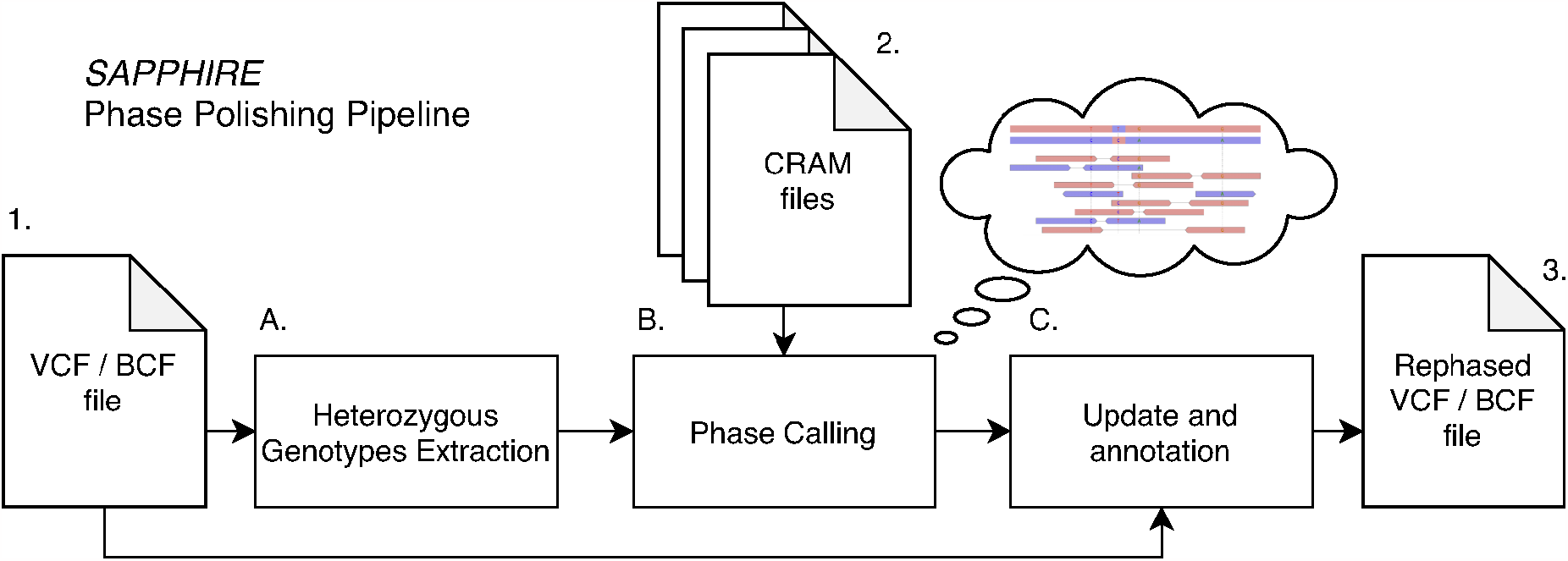
SAPPHIRE phase polishing pipeline. From an input VCF/BCF file **1** (e.g., whole chromosome) heterozygous genotypes are extracted **A** to be rephased alongside their closest neighboring heterozygous genotypes. The phase calling stage **B** accesses the whole-genome sequencing files (CRAM files **2**) to call the phase of extracted heterozygous genotypes in relation to their neighbors. Finally the original VCF/BCF file 1 is updated and annotated **C** to generate the final rephased VCF/BCF file **3**.

## Data

We used the whole-genome sequencing data available on the UK Biobank RAP as pVCF files that were called with GraphTyper [18]. We used two different releases of the UK Biobank WGS data: (a) the first release of 150,119 individuals, to compare the SAPPHIRE method to the original phasing with the SHAPEIT5 method [3], and (b) the newest release of 200,031 individuals. For the release of the 150,119 individuals with WGS data available, we performed quality control of this data as described previously [3, 19]. Briefly, we (i) used BCFtools [20] to split multi-allelic variants into bi-allelic variants, (ii) removed SNPs and indels having more than 10 percent of missing genotypes, Hardy-Weinberg p-value *<* 10^*−*30^ and an excess of heterozygosity less than 0.5 or greater than 1.5. Additionally, we (iii) filtered out variants with AAscore *<* 0.5, and (iv) kept only variants with the PASS FILTER tag. In total, this filtering retained 603,925,301 variant sites across 150,119 individuals. For our analysis, we kept only individuals being also genotyped with the UK Biobank Axiom array. Additionally, since we used white British trios and duos (*N* = 31 and 432 individuals, respectively) to assess the accuracy of the method as previously described [3], we removed parental genomes from the data. This resulted in a total of 147,754 individuals. The quality control of the second UK Biobank WGS data release (i.e., 200,031 individuals) has been performed as part of the official release of the phased data by the authors of SHAPEIT5 [3]. Although very similar to the previous quality control, it has been optimized for phasing accuracy and only variants with AAscore *>* 0.8 were kept. In this release, family phasing was applied in order to use parental genomes to phase the offspring (*N* = 93 trios and *N* = 915 duos).

### Extraction of heterozygous genotype data for polishing

First, for each sample, all heterozygous genotypes phased by SHAPEIT5 with a phasing confidence score (PP field in VCF) below 0.99 (99% confidence) were extracted alongside their closest neighboring heterozygous genotypes (up to two before and two after, the closest neighboring heterozygous variants are different for each sample) regardless of their phasing confidence score. The extracted heterozygous genotypes and their neighbors are stored in a binary format and the variant loci information is kept in BCF format (without any sample). Splitting the information this way allows for faster random access [21]. The generated files are then passed along to the next stage, the phase caller, which will query whole-genome sequencing reads to verify the phase calls.

### Phase calling and polishing of extracted sites

Extracted heterozygous sites are filtered on variant type and only single nucleotide variants (SNVs) are processed, as shown in Fig 5. For each site a pileup of sequencing reads is done through HTSLIB [22] similarly to the pileup function of BCFTools [20]. For each site that has a phasing confidence score below 0.99, evidence from sequencing reads is queried and the number of reads that confirm or invalidate the phase of the site is counted. If there is enough evidence (at least one high quality mapped sequencing read and no contradicting reads) to show a phase error, the phase is flipped as shown in Fig 1B. Evidence, the number of reads that confirm the new phase, is recorded. For all sites that are phased correctly, the number of reads is also recorded in the phase confidence score to show that the phase has been validated by sequencing reads.

### Read quality filtering

In order to avoid lowering the quality of the input data sets to be polished, we applied a very strict filter on which sequencing reads are allowed to be used as evidence. A sequencing read-pair would be rejected by the SAPPHIRE algorithm if : Mapping quality score (MAPQ) is below 50, or one of the sequencing reads in the pair is unmapped, or a read is unpaired, or the read pair has incorrect orientation (e.g., both reads on the forward strand), or a read has the duplicated flag set. This way only very high quality aligned reads remain, which at one side results in less genotypes getting their phase validated or switched, but on the other side, newly validated or switched genotypes are of very high confidence. Adjustment of this read quality filtering could allow it to phase polish more genotypes, evaluation of a more relaxed filtering is left for future works.

### Scatter-Gather parallelism

In order to reduce the time needed to process all samples on the UK Biobank RAP, the extracted data is split in batches of samples, e.g., 1,000 samples, 5,000 samples., 10,000 samples. Because all samples are processed individually with their respective whole-genome sequencing data, this process can be multi-threaded up to a certain degree dictated by the access bandwidth to the whole-genome sequencing files per node. For example we empirically found out that running a dozen threads on a four CPU node would give the most efficient node usage. The number of threads being higher than the CPU count is so that enough threads are working while other threads are waiting for data to arrive over the network.

### Accessing a large number of sequencing files

Since the whole-genome sequencing read files (in CRAM format [23, 24]) are on average more than 17 GB per sample resulting in 3,669,126 GB for all 200,031 samples, naive approaches are not tractable. The jobs running SAPPHIRE were split per chromosome in batches of 1,000, 5,000, 10,000 samples based on chromosome size (details in S1 Table). Each job therefore requires access to at least 1,000 CRAM files. Jobs on UK Biobank RAP are run in compute nodes (backed by Amazon AWS instances) and can access CRAM files in two ways, first, by downloading the file, then processing it, second, by mounting the UK Biobank data as a read-only network file system. The first approach is not viable as it would first require a large amount of storage on the compute node and would spend a long time moving data before processing (1,000 samples ≈ 17 TB of data). Therefore CRAM files were accessed over a network file system. HTSLIB, which allows for random access in indexed CRAM files and this enables the processing of a large number of samples on a single node. The main bottleneck becomes network speed, measured at about 50 MB/s on average. Thanks to the random access features of the CRAM format and HTSLIB, only the required parts of the CRAM files are accessed. The CRAM format also provides a checksum for each of its internal data records, allowing it to detect errors in network transfers and act accordingly.

### Update and annotation of the original VCF/BCF file

Finally, the original VCF/BCF file is updated with the new phase and phase confidence scores. This stage also checks that no errors have been introduced (e.g., a heterozygous genotype call becoming a homozygous call or that a high confidence heterozygous site would have been rephased). The new phase confidence score is defined as follows : old confidence score plus number of reads that confirm the phase plus one. So all genotypes with a score over 1.0 are polished genotypes. The original phase confidence can be extracted by looking at the fractional part, with the edge case of 1.0 which is a phase confidence of 1.0, which we don’t rephase.

### Validation of haplotype estimates

To validate the haplotype estimates and compare the accuracy of methods, we used parent-offspring trios and duos that we identified using the kinship estimate and the IBS0 provided in the UK Biobank Axiom array release. Specifically, we infer parent-offspring relationships for any pair of individuals having kinship coefficient lower than 0.3553 and greater than 0.1767, and IBS0 lower than 0.0012 [1, 3, 25]. We then used the age to infer the direction of the relation-ship. Additionally, we kept only white British individuals for which the ancestry was confirmed by Principal Component Analysis (PCA). It resulted in 31 trios and 432 duos across the 150,019 individuals, and 93 trios and 915 duos across the 200,031 individuals. The validation process consisted in phasing the data set excluding parental genomes, and using parental genomes to assess how close are the estimate from the true parental haplotypes, measured as a switch error rate (SER) which is defined as the fraction of successive pairs of heterozygous genotypes being correctly phased [3]. Alternatively, we used genotype imputation to assess the accuracy of our method regardless of family information. Notably, this allows us to validate our haplotypes for the second data release in which parental genomes were kept. For this purpose, we selected 1,000 individuals of white British ancestry that are unrelated to any other sample in the data set and for which SNP array data is available, as traditionally done in imputation experiments [19]. We removed these individuals from the phased data sets, constituting reference panels of 146,754 and 199,031 individuals. We used the UK Biobank Axiom array data as input for genotype imputation. We quality control the input data using the UKB SNPs and samples QC file (UKB Resource 531) as previously described [3]. We filtered out (i) variants that were not kept in the official phasing of the Axiom array data [1] and (ii) variants with a difference greater than 0.1 in allele frequency between the Axiom array data and the reference panel [3]. We used IMPUTE5 [26] for genotype imputation of the SNP array data and the concordance tool of GLIMPSE2 [19] to assess imputation accuracy.

### Compute nodes and cost

The heterozygous genotype extraction (A in Fig 9) can be run on a single node per chromosome. For all samples the compute node was chosen on storage size due to the size of the input VCF files. The phase calling and polishing (B in Fig 9) is done per batch of 1,000-10,000 samples per node depending on chromosome size. Finally the update of each VCF file is run on a single node per chromosome (C in Fig 9). The total compute time and compute cost (time × price per node) is reported in S1 Table. In order to polish the phase of all 200,031 samples in the UK Biobank the total cost was £570 with ≈ 60% of the cost spent on the actual phase calling with WGS data in the SAPPHIRE method (B in Fig 9). Compute nodes for stage A were chosen on lowest cost nodes that would provide enough solid state storage to store the input VCF file. Compute nodes for stage B were selected empirically to provide the best price-performance with relation to the network speed of the network attached storage that holds all the WGS data. Finally for stage C the nodes were chosen based on the storage for both input and output files. Note: There are two priorities *on-spot* (low priority) and *on-demand* (high priority) for nodes on the UK Biobank RAP and the difference in cost is about five-fold. Low priority jobs can be interrupted if there are no nodes left for high priority jobs. If this happens, the interrupted job will be relaunched with high priority, resulting in higher cost. This explains the variability of costs reported in S1 Table. The final cost of running the SAPPHIRE method pipeline per sample on the 200k UK Biobank release was £570 / 200,031 *≈* £0.00285, less than a single penny per sample.

### Cost estimation of read based phasing

Alternatively to phase polishing statistically phased haplotypes it would be possible to apply read based phasing first, e.g., with WhatsHap [5] to generate phase sets and then in a second pass apply a statistical method that takes into account phase sets, e.g., SHAPEIT4 [6]. Running WhatsHap for a single sample on chromosome 22 with whole-genome sequencing took 14min43 on a UKB RAP mem3_ssd2_v2_x8 node. WhatsHap reported: 157s reading CRAM, 85s parsing VCF, 25s selecting reads, 50s phasing, 90s writing VCF, and 476s uncategorized. The maximum memory usage was 10.8 GB. This benchmark was run with a VCF already sub-sampled to a single sample because opening the 200,031 sample VCF file with WhatsHap to phase a single sample did run out of memory on the 64 GB node. Note that extracting a single sample from the 200,031 sample VCF file took 262 minutes with BCFTools on a UKB RAP `mem2_ssd2_v2_x2` node with files stored locally on the SSD. We can estimate running WhatsHap alone for 200,031 samples will take about 50,000 CPU hours (200, 031 × 15min). Given the memory requirements per sample we can run up to five processes in parallel on a single 64 GB node reducing the effective time to 10,000 hours. We can compare this to the SAPPHIRE method runtimes in S1 Table. It takes 9h19min to extract the positions to phase polish for all samples on chromosome 22. The phase polishing itself takes 7h14 on 21 nodes, so 153 hours, followed by a final update of the VCF with all samples, which takes 24h28min. Therefore the total time is 187 hours to phase polish 200,031 samples from VCF to VCF. If we compare the 10,000 hours running WhatsHap to 187 hours of phase polishing we can say that running WhatsHap would take at least 50 × more time than SAPPHIRE and this does not even include time to extract the samples from the VCF, nor applying the phase sets to another method, nor consolidating the results. The cost of SAPPHIRE for chromosome 22 was £4.47, the estimated cost for WhatsHap on chromosome 22 is £528 (10,000 hours at a £0.0528 per hour rate, the lowest rate for a 64 GB instance in the UKB RAP). The estimated cost is at least 100 × more than SAPPHIRE. The SAPPHIRE method efficiency comes from several factors: First, it only considers variants with low phasing confidence (or low MAF as a proxy) where WhatsHap considers all heterozygous variants. Second, the extraction step of the SAPPHIRE method converts the input VCF to a sparse binary format that allows extremely rapid parsing of variants. Third, extraction is done directly as a stream to a file, therefore memory usage of the SAPPHIRE pipeline is very low and the extraction can be run on nodes with very limited memory (e.g., 8 GB) whereas WhatsHap failed to load a single sample from the 200,031 samples VCF on a 64 GB instance. Therefore, we can conclude that a lower bound for the cost of using a read based phasing method would be at least 100 × more expensive than phase polishing with SAPPHIRE. Phase polishing all autosomes of 200,031 samples of the UKB WGS data set with SAPPHIRE did cost £570, an estimation for a full read based phasing method would be at least £57,000.

## Software availability

The software developed for the SAPPHIRE method is open-source and available under non-restrictive license. The software is combined as a toolkit and can be built and run on the UK Biobank RAP. The software is available at https://github.com/rwk-unil/pp.

## Author Contributions

R.W. and O.D. designed the study. R.W. and O.D. developed the algorithms. R.W. wrote the software. R.J.H. performed the statistical phasing experiments. R.W. performed all other experiments. I.X. provided interpretation of the results. O.D. and Y.T. supervised the project. All authors reviewed the final manuscript.

## Supporting information

**S1 Table.**
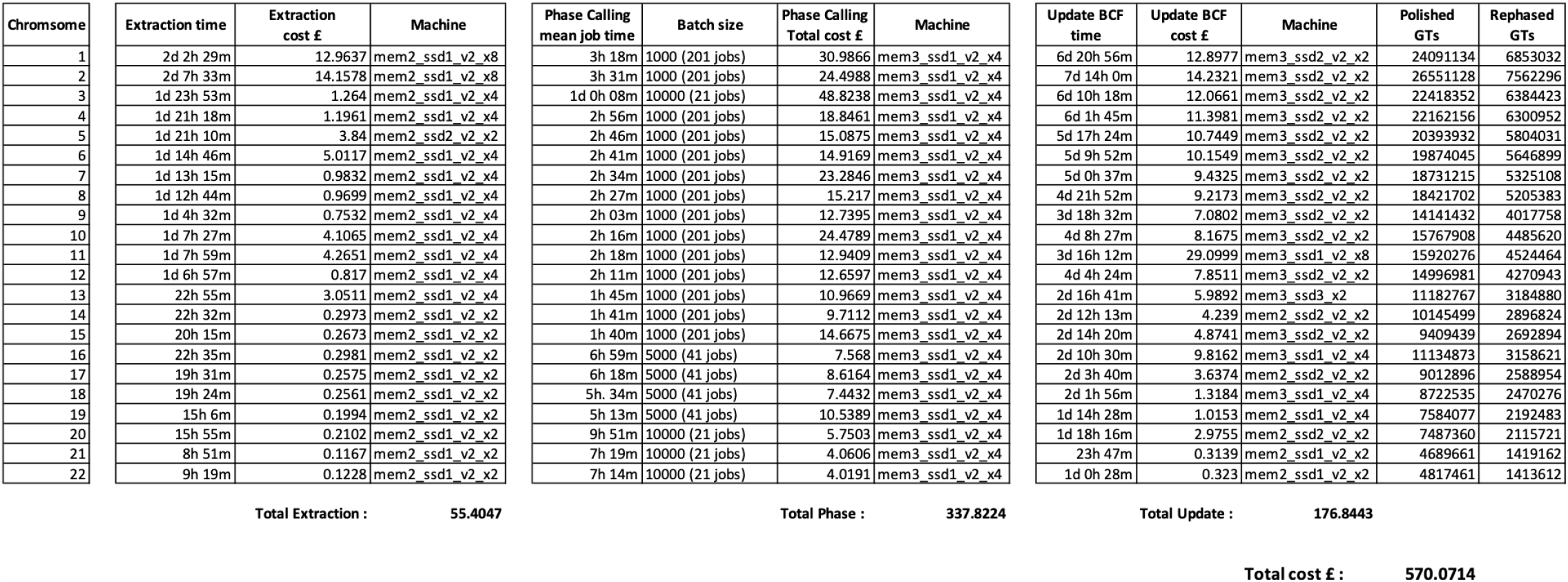
Cost of the SAPPHIRE Phase Polishing pipeline on the 200k UK Biobank release. The cost for all three steps of the phase polishing pipeline for chromosomes 1-22 of the UK Biobank 200k release. The final number of polished genotypes (GTs) and rephased GTs is given for all chromosomes. Notes: Five-fold increases in cost are for jobs that were interrupted and had to be relaunched with higher priority, for example extraction on chromosome 6 and 7 almost take the same time on the same machine but the job of chromosome 6 was interrupted and relaunched at a higher priority (and cost). Chromosome 3 had the phase calling jobs split into batches of 10,000 samples as a test, which showed that it was better to split big chromosomes in more batches of smaller sample size. This has two advantages, first, the wall clock time is reduced and second, the cost is reduced because with the large sample size, the jobs had to run for a long time and were interrupted and had to be relaunched with high priority which increased the cost.

## Acknowledgments

The benchmarks on the UK Biobank data have been conducted using the UK Biobank Resource under Application Number 66995. This work has been supported by the School of Engineering and Management Vaud (HEIG-VD). O.D. was supported by SNF grant number: SNSF-PP00P3 176977.

## Notes

### Competing Interest Statement

The authors have declared no competing interest.

